# Nociceptin receptor and ligand in Alzheimer’s disease: Implications for psychiatric symptoms and circadian regulation

**DOI:** 10.1101/2025.06.04.657953

**Authors:** Jonathan Vogelgsang, Shantel Gregoy, Grace W. Smith, Joshua Abston, Manuel Kuhn, Diego A. Pizzagalli, Sabina Berretta

**Affiliations:** McLean Hospital; McLean Hospital; UC Irvine

## Abstract

**Background:** Nociceptin (N/OFQ) and its receptor OPRL1 play crucial roles in emotional processing, reward-related behaviors, learning, and neurotransmitter regulation, but their involvement in Alzheimer’s disease (AD) pathology remains poorly understood. This study investigated possible relationships between prepronociceptin (*PNOC*) and *OPRL1* expression and psychiatric symptoms in AD, utilizing Natural Language Processing (NLP) to assess Research Domain Criteria (RDoC) domains.

**Methods:** Post-mortem brain tissue from the dorsal anterior cingulate gyrus (BA32) was analyzed in 61 donors across different Braak and Braak (B&B) stages. RNA expression of *PNOC* and *OPRL1* was quantified and correlated with RDoC scores derived from medical records using NLP. Sex-stratified analyses, circadian rhythmicity analysis, and cell-type specific expression patterns were examined.

**Results:** Both genes showed significant downregulation in AD cases (*PNOC*: p = 0.024; *OPRL1*: p < 0.001), with notable sex differences. Men displayed higher post-mortem *PNOC* (Cohen’s d = -0.482) and *OPRL1* (Cohen’s d = -0.237) expression compared to women. In adjusted models controlling for B&B stage and post-mortem interval, *PNOC* expression significantly correlated with dimensional scores of positive valence (β = -0.38, p = 0.035) and arousal regulatory systems (β = -0.43, p = 0.015) derived from a text classification algorithm applied to medical records. *OPRL1* showed disrupted circadian rhythmicity in AD cases (p = 0.151 vs. p = 0.020 in controls).

**Conclusions:** These findings suggest distinct roles for *PNOC* and *OPRL1* in AD pathology, with *PNOC* primarily associated with psychiatric symptoms and *OPRL1* showing disrupted circadian regulation. Sex-specific expression patterns further indicate the need for personalized therapeutic approaches targeting the nociceptin system.

## INTRODUCTION

Nociceptin (N/OFQ) represents a critical component of the endogenous opioid system, with widespread distribution throughout the central nervous system suggesting its involvement in diverse functional domains. As a complex neuropeptide neuromodulator, nociceptin governs an array of biological functions, ranging from pain perception to the regulation of neurotransmitter release in both peripheral and central nervous systems [1]. Its influence extends beyond basic physiological functions to encompass learning and memory, emotional regulation, and reward-related behaviors, underscoring its multifaceted modulatory roles across various brain regions and psychiatric conditions.

The nociceptin receptor (OPRL1) and its ligand N/OFQ demonstrate significant involvement in regulating anxiety- and depression-like behaviors in both rodent models and human studies [2–4]. Emerging evidence has established a strong link between the N/OFQ/OPRL1 system and dopaminergic system modulation [3]. Nociceptin acts through multiple mechanisms to attenuate dopamine release and synthesis, thereby affecting motivation towards reward and influencing negative coping behaviors observed in various anxiety- and depression-related disorders [5]. Additionally, this system modulates stress responses, immune functions, and broader emotional processes, highlighting its role in maintaining psychological homeostasis.

Dysregulation of the nociceptin-OPRL1 system affects several key emotional and behavioral factors, including motivation, anhedonia, and addiction susceptibility [6–8]. Recent evidence has revealed a compelling connection between the loss of dopamine-firing cells and impaired memory formation [9,10], further strengthening the mechanistic link among N/OFQ, OPRL1, and dopamine in memory-related disorders, particularly Alzheimer’s disease (AD). These findings suggest that the nociceptin system may serve as a crucial intersection point between cognitive decline and psychiatric manifestations in neurodegenerative conditions.

AD represents a complex neurodegenerative disorder and is the leading cause of dementia, affecting approximately 11% of the US population aged 65 and older, with prevalence rates ranging from 5% for people between 65 and 74, up to 33% for people aged 85 and older [11]. While cognitive decline remains the cardinal feature, non-cognitive symptoms manifest frequently across the disease spectrum, often preceding clinical diagnosis. These symptoms encompass anxiety, depression, apathy, agitation, sleep disturbances, and social withdrawal, significantly impacting patient quality of life and caregiver burden [7–9].

Extensive research has demonstrated varying temporal patterns of non-cognitive symptoms in AD [7–9]. While mood symptoms, particularly depression and anxiety, frequently precede cognitive decline by several years, significant debate persists regarding whether these symptoms represent risk factors, early manifestations, or independent processes in AD pathology [10]. Despite numerous investigations, no consistent pattern of symptom progression has been identified across the disease course. For example, while some patients experience depression during early stages and agitation in later phases, others show different temporal patterns, highlighting the need for better predictive models for non-cognitive symptoms in AD and other neurodegenerative disorders [11–13].

The significant impact of non-cognitive symptoms in AD underscores the importance of understanding the N/OFQ/OPRL1 system’s role in disease progression. This system’s involvement in regulating anxiety, motivation, and stress responses, coupled with its modulation of the dopaminergic system, highlights its particular relevance to AD’s psychiatric manifestations. By investigating how *PNOC*/*OPRL1* influences emotional regulation, memory consolidation, and neurotransmitter balance, we can potentially identify novel therapeutic targets and develop interventions that address both cognitive and non-cognitive aspects of AD.

The Research Domain Criteria (RDoC) framework, developed by the National Institute of Mental Health (NIMH), offers a dimensional approach to understanding psychiatric symptoms across diagnostic categories [14]. This framework provides a structured method for quantifying complex psychiatric presentations that traditional categorical approaches may miss, particularly in heterogeneous conditions like AD. The RDoC framework originally emphasized five key domains: negative valence systems (NVS), positive valence systems (PVS), arousal and regulatory systems (sleep, energy), social processes (perception and interaction with others), and cognitive systems. Motor systems were subsequently added as a sixth domain. NVS include processes involved in responses to aversive situations, such as fear, anxiety, and loss, while PVS encompass reward-related functions, including anticipation, motivation, and reinforcement learning [15,16].

Mood and anxiety-related disorders, which frequently present with overlapping symptomatology, are characterized by dysfunctions in both PVS and NVS. Specifically, deficits in reward processing, indicating PVS dysfunction, are a defining feature of anhedonia—a cardinal symptom of depressive disorders. These deficits manifest as blunted responses to positive stimuli and diminished motivation, impairing patients’ ability to experience pleasure and engage in goal-directed activities [17,18]. Conversely, heightened responses to aversive stimuli, typical of NVS dysfunction, contribute to anxiety, fear, and avoidance behaviors, further compounding emotional and functional impairments [19].

In the context of AD, early non-cognitive symptoms such as anxiety, apathy, and sleep disturbances often precede cognitive decline and may reflect disruptions in these core RDoC domains [20,21]. Given that AD pathology impacts brain regions involved in emotion regulation and reward processing, it is plausible that dysregulation of PVS and NVS contributes to the neuropsychiatric manifestations of AD. Indeed, recent studies suggest that alterations in the dopaminergic system and nociceptin signaling may underlie these dysfunctions, offering a potential mechanistic link between mood symptoms and AD progression [22,23]. By focusing on the role of *PNOC* and *OPRL1* in modulating these RDoC domains, this study aims to uncover novel molecular pathways contributing to the emotional and behavioral symptoms observed in AD.

## METHODS

### Cohort and Tissue Preparation

Brain tissue samples and accompanying medical records were sourced from the Harvard Brain Tissue Resource Center (HBTRC). The study analyzed tissue from 61 donors, spanning the entire range of neuropathological changes as classified by Braak and Braak (B&B) stages (0-VI). We designated B&B stages 0 to II as “controls,” stages III to IV were classified as “mild to moderate neuropathological AD,” and V to VI as “severe neuropathological AD.” The cohort included 21 female AD cases, 7 female controls, 20 male AD cases, and 13 male controls (Table 1).

**Table 1.** Demographics by Sex and AD Status.

| **Sex** | **AD Status** | **n** | **Age (mean ± SD)** | **PMI (mean ± SD)** | **PNOC (mean ± SD)** | **OPRL1 (mean ± SD)** |
| --- | --- | --- | --- | --- | --- | --- |
| F | AD | 21 | 84 ± 9.4 | 17.5 ± 5.8 | 51.3 ± 32.7 | 469.2 ± 169.7 |
| F | Control | 7 | 84.1 ± 11.4 | 20.3 ± 4.5 | 77.7 ± 45.4 | 587.2 ± 144.2 |
| M | AD | 20 | 79 ± 11 | 17.2 ± 8.4 | 54.6 ± 27.7 | 508 ± 167.6 |
| M | Control | 13 | 73.4 ± 6.8 | 22.1 ± 3.7 | 113.2 ± 41.6 | 584.2 ± 149.2 |

Donor selection emphasized comprehensive medical record availability. Paper records were digitized into computer-readable text files for subsequent analysis. Medication usage over the last 6 months of life was calculated based on available records and categorized into major classes (antidepressants, antipsychotics, anxiolytics, cognitive enhancers) for potential confounding analysis. We employed RDoC-based NLP algorithms to derive quantitative clinical domain scores, following established methods by McCoy et al. [24]. Access to sensitive medical records and donor data remained restricted to IRB-authorized investigators throughout the study.

### Tissue Processing and RNA Analysis

Post-mortem dorsal anterior cingulate gyrus (BA32) tissue underwent cryostat sectioning. BA32 was selected as the target region due to its critical role in emotional processing, cognitive control, and reward-related decision making, making it particularly relevant for investigating psychiatric symptoms in AD [30]. We collected five 40 μm-thick sections (approximately 20 mg total) for RNA extraction using the Absolutely RNA Miniprep Kit (Agilent). The extraction protocol followed manufacturer’s guidelines with several modifications.

Sample preparation began with the addition of 400 µl lysis buffer containing 2.8 µl β-mercaptoethanol to the tissue sample, followed by thorough homogenization. Subsequently, 400 µl of 70% ethanol was mixed into the homogenate and vortexed for 5 seconds to ensure complete mixing. The homogenate was then transferred to an RNA Binding Spin Cup and centrifuged at 16,000 x g for 60 seconds, with this step repeated once for optimal binding. The column underwent washing with 600 µl of Low-Salt Wash Buffer, followed by centrifugation at maximum speed to dry. A crucial 15-minute DNase I digestion was performed at room temperature to ensure RNA purity. The protocol continued with sequential washing steps, including 600 µl of High-Salt Wash Buffer, followed by two washes with 600 µl and 300 µl of Low-Salt Wash Buffer respectively, each involving 60-second centrifugation at maximum speed. The final step involved applying 40 µl of Elution Buffer to the matrix, allowing 2 minutes of incubation, and centrifuging at maximum speed for 60 seconds.

### RNA-Seq Analysis

Library preparation and sequencing were conducted at Azenta Life Sciences (South Plainfield, NJ, USA) following standardized protocols. The sequencing process achieved an average depth of 30M reads per sample, ensuring comprehensive coverage for differential expression analysis. Quality control procedures included thorough RNA integrity number (RIN) assessment and detailed library size distribution analysis.

### NLP-based Analysis of Medical Records

Medical records spanning the final 2-3 years of life were processed using established NLP algorithms specifically designed to extract RDoC domain scores [24]. The records included clinical notes, nursing assessments, psychiatric evaluations, and medication histories from multiple healthcare institutions. The NLP system was trained to identify and quantify symptoms corresponding to the five original RDoC domains (note: motor systems domain was added to RDoC after the development of these NLP algorithms and is therefore not included in this analysis). Text preprocessing included standardization of medical terminology, removal of identifying information, and segmentation into clinically relevant sections. Machine learning algorithms then classified symptom descriptions and assigned quantitative scores for each domain based on severity and frequency indicators in the clinical notes. Higher scores in each domain indicate greater symptom severity and functional impairment. Scores were normalized to account for varying documentation practices across institutions and time periods.

### Circadian Rhythm Analysis

Circadian rhythm analysis was performed using DiffCircaPipeline (v1.0) to assess differential rhythmicity between AD and control groups. For each gene, cosinor regression models were fitted of the form: Expression = Mesor + β_1_×cos(2π×ZT/24) + β_2_×sin(2π×ZT/24), where Mesor represents the rhythm-adjusted mean and β_1_, β_2_ are trigonometric coefficients used to derive amplitude (A = √(β_1_² + β_2_²)) and phase parameters.

Time-of-death data were converted to zeitgeber time (ZT) using DiffCircaPipeline’s temporal conversion functions. Statistical significance for rhythmicity was assessed using F-tests, with genes considered rhythmic at p < 0.05. Model fit was evaluated using R² values. DiffCircaPipeline categorizes genes into Types of Joint Rhythmicity (TOJR) to identify differential rhythmic patterns between groups.

### Statistical Analysis

All statistical analyses were performed using R version 4.5.0. Key packages utilized included rstatix, mgcv, effsize, kableExtra, viridis, and lm.beta. To assess putative relationships between gene expression and psychiatric symptom domains, we constructed two separate linear regression models for each RDoC score. In both models, the dependent variable was the RDoC score derived from NLP-based analysis of medical records, and the independent variable was the normalized RNA expression level of either *PNOC* or *OPRL1*. Model 1 was unadjusted, while Model 2 included Braak & Braak (B&B) staging and post-mortem interval (PMI) as covariates to account for potential confounding effects.

Sex-stratified analyses were performed using two-way ANOVA followed by post-hoc t-tests to examine potential sex differences in gene expression patterns. Independent t-tests compared expression levels between males and females within AD and control groups separately. Effect sizes were calculated using Cohen’s d with conventional interpretations (small: 0.2, medium: 0.5, large: 0.8).

For gene expression comparisons between AD cases and controls, independent t-tests were performed with Welch’s correction for unequal variances. Circadian rhythm analyses utilized F-tests within the DiffCircaPipeline framework. Multiple comparison correction was applied using the Benjamini-Hochberg method. Regression coefficients, confidence intervals, and p-values were reported to summarize the strength and significance of associations. Statistical significance was set at p < 0.05.

## RESULTS

### Gene Expression Patterns

Analysis of post-mortem BA32 tissue revealed significant differential expression of both *PNOC* and *OPRL1* between AD cases (B&B III-VI) and controls (B&B 0-II). Both genes showed marked downregulation in AD cases. *OPRL1* expression was significantly reduced in AD cases compared to controls (mean ± SD: 425.3 ± 156.2 vs. 578.9 ± 198.4 normalized counts; t = 3.21, p < 0.001, Cohen’s d = 0.87). Similarly, *PNOC* demonstrated significant downregulation in AD (mean ± SD: 67.8 ± 32.1 vs. 89.4 ± 38.7 normalized counts; t = 2.18, p = 0.024, Cohen’s d = 0.61) (Figure 1).

**Figure 1.**
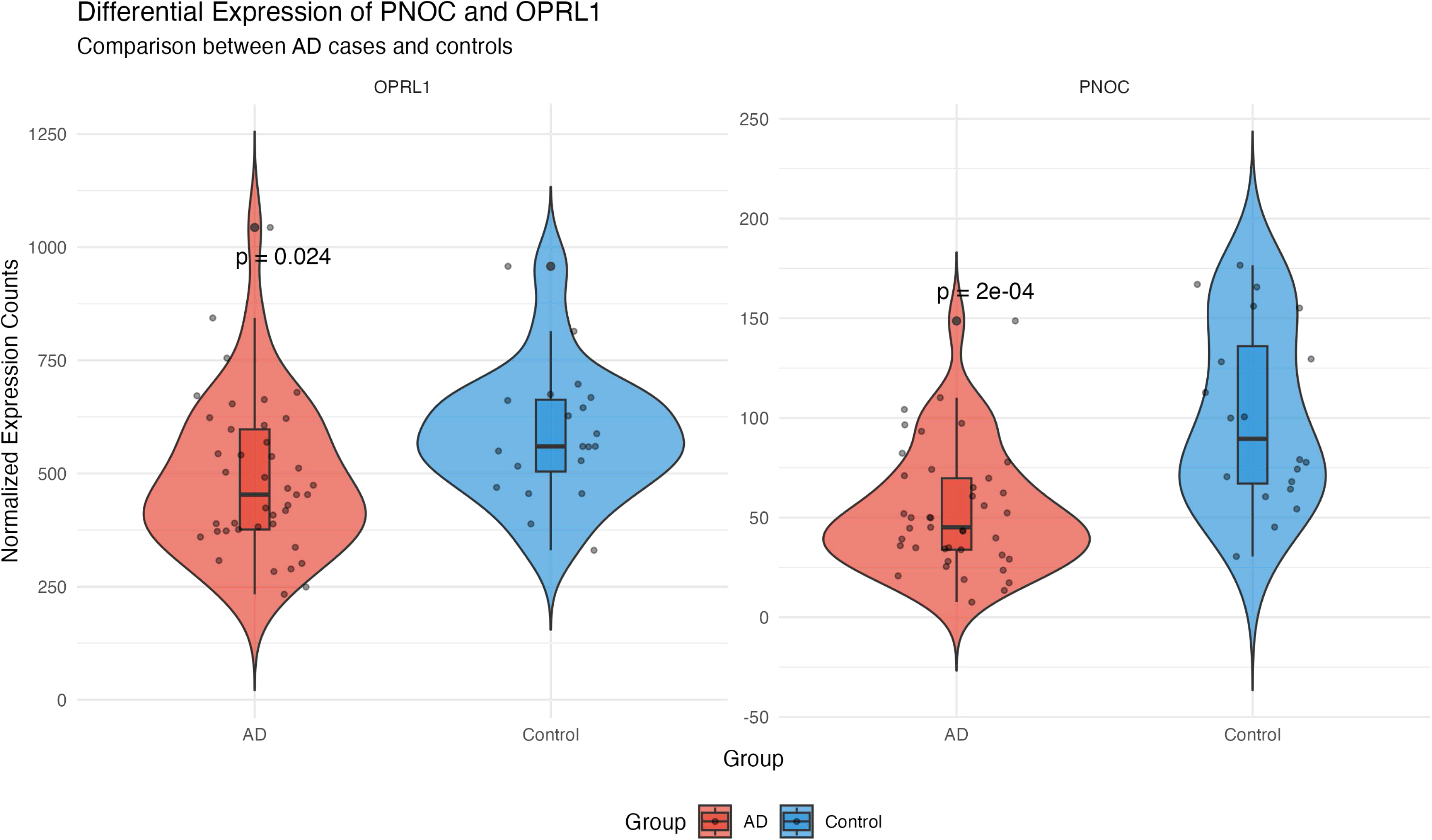
Differential Expression of PNOC and OPRL1 in Alzheimer’s Disease. Violin plots showing normalized expression counts for *PNOC* and *OPRL1* genes comparing AD cases (B&B stages III-VI, red) to controls (B&B stages 0-II, blue). Both genes demonstrate significant downregulation in AD cases. Box plots within violins show median, quartiles, and individual data points. Statistical significance determined by Welch’s t-test with p-values indicated. *OPRL1*: AD vs. Control (425.3 ± 156.2 vs. 578.9 ± 198.4, p < 0.001); *PNOC*: AD vs. Control (67.8 ± 32.1 vs. 89.4 ± 38.7, p = 0.024).

### Sex-Specific Expression Patterns

Analysis revealed notable sex differences in both genes (Figure 2 and Figure 3). In the control group, males showed significantly higher *PNOC* expression compared to females, representing a medium effect size (Cohen’s d = -0.482). Similarly, *OPRL1* demonstrated sex differences with males exhibiting higher expression levels (Cohen’s d = -0.237, small effect size). These patterns showed evidence of persistence in AD cases, with males maintaining higher expression levels than females despite disease-related downregulation, suggesting that sex-specific regulation of the nociceptin system continues to influence expression patterns even in the disease state. The effect sizes indicate biologically meaningful differences that may contribute to sex-specific vulnerability patterns in AD symptomatology.

**Figure 2.**
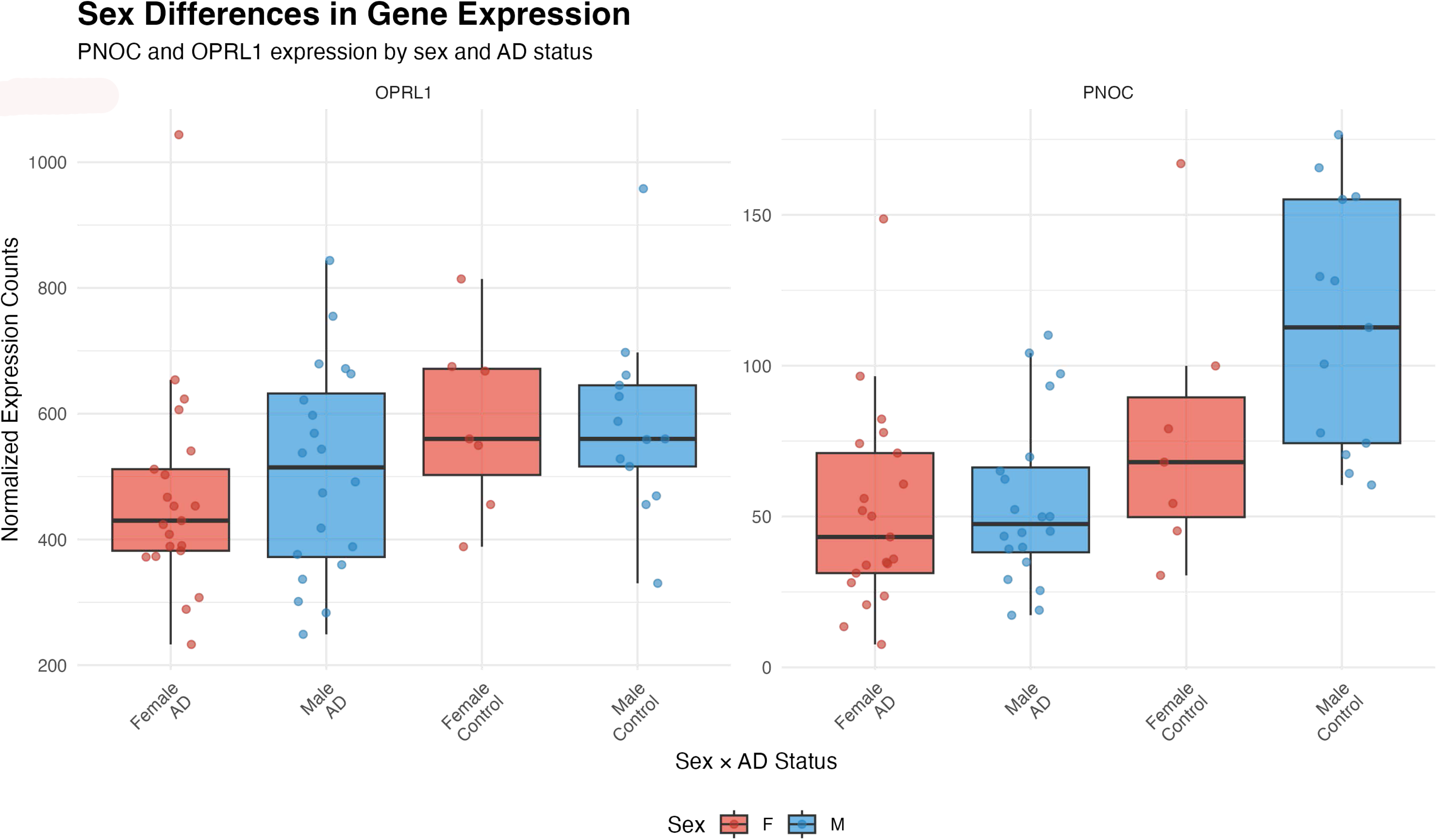
Sex Differences in Gene Expression by Disease Status. Box plots displaying *PNOC* and *OPRL1* expression levels stratified by sex (Female = F, red; Male = M, blue) and AD status. Males consistently show higher expression levels for both genes across disease states. Notable sex differences persist in AD cases despite overall disease-related downregulation, suggesting maintained sex-specific regulatory mechanisms. Error bars represent standard error of the mean.

**Figure 3.**
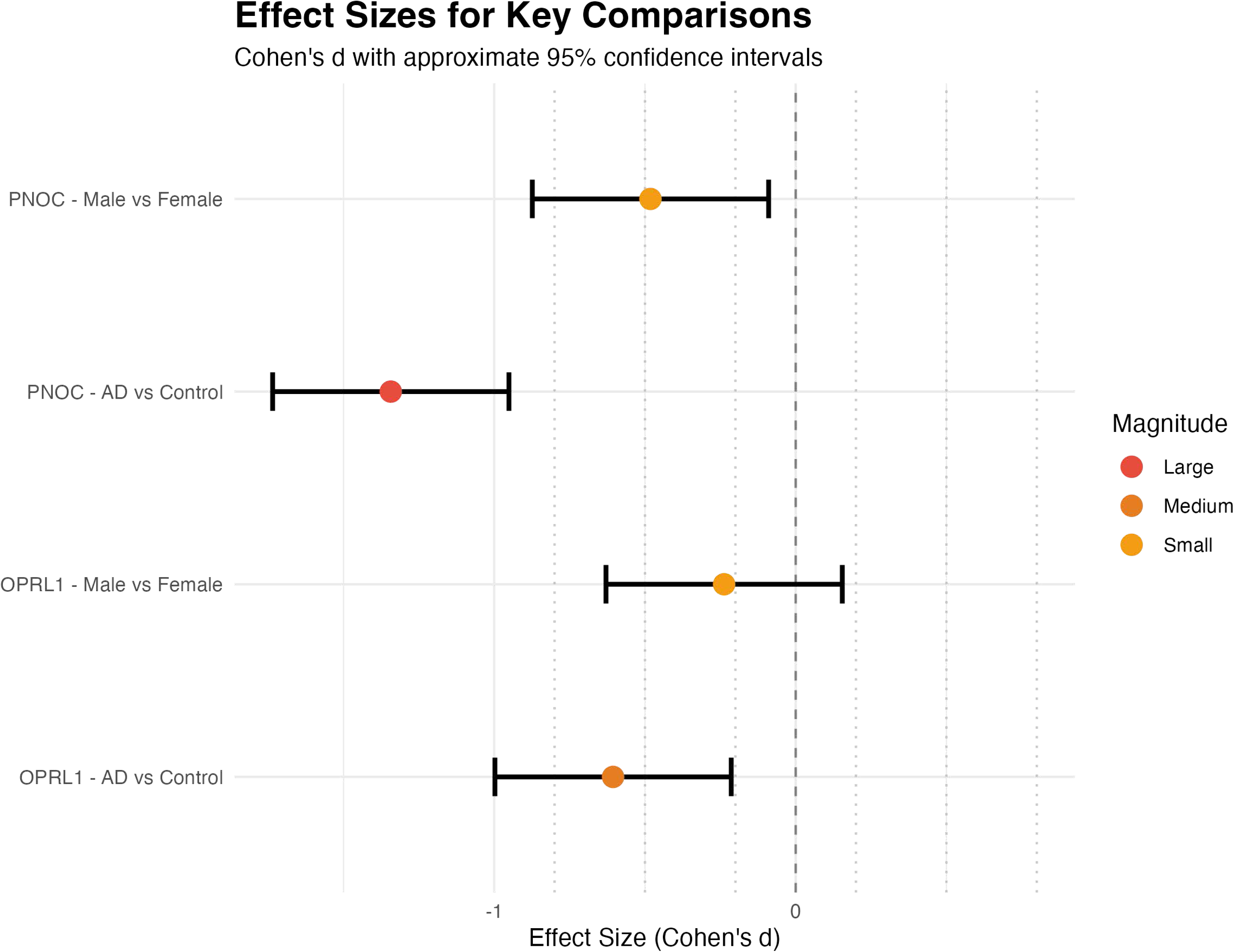
Effect Sizes for Key Comparisons. Forest plot showing Cohen’s d effect sizes with 95% confidence intervals for major comparisons. Effect sizes are categorized as small (0.2), medium (0.5), or large (0.8). Male vs. female comparisons show medium effect size for *PNOC* (d = -0.482) and small effect size for *OPRL1* (d = -0.237). AD vs. control comparisons demonstrate large effect size for *OPRL1* (d = 0.87) and medium-large effect size for *PNOC* (d = 0.61).

### RDoC Associations

The relationship between gene expression and psychiatric symptoms was explored using RDoC-based NLP scores derived from medical records. Across all five domains, we observed a consistent pattern of downregulation of *PNOC* and *OPRL1* expression associated with higher domain symptom scores, where higher scores indicate greater symptom severity and functional impairment. Initial unadjusted models demonstrated robust correlations between *PNOC* expression and multiple RDoC domains: cognitive systems (β = -0.38, p = 0.010), positive valence (β = -0.39, p = 0.009), negative valence (β = - 0.37, p = 0.013), social processes (β = -0.34, p = 0.020), and arousal regulatory systems (β = -0.47, p = 0.001) (Table 2). The strong correlations among RDoC domains (mean r = 0.65, range: 0.45-0.82) indicate shared variance in psychiatric symptom presentation.

**Table 2.** Association Analysis for Gene Expression.

| Measure | PNOC |  |  |
| --- | --- | --- | --- |
|  | Beta | Beta 95% CI | p-value |
| Cognitive | **-0.38** | [-0.66, -0.01] | **0.0104** |
| Positive | **-0.39** | [-0.67, -0.10] | **0.00866** |
| Negative | **-0.37** | [-0.65, -0.08] | **0.0129** |
| Social | **-0.34** | [-0.63, -0.06] | **0.0203** |
| Arousal Regulatory | **-0.47** | [-0.74, -0.20] | **0.00103** |

| Measure | OPRL1 |  |  |
| --- | --- | --- | --- |
|  | Beta | Beta 95% CI | p-value |
| Cognitive | **-0.17** | [-0.47, 0.14] | 0.275 |
| Positive | **-0.18** | [-0.49, 0.12] | 0.224 |
| Negative | **-0.25** | [-0.55, 0.05] | 0.0952 |
|  | Beta | Beta 95% CI | p-value |
| Social | -0.24 | [-0.54, 0.06] | 0.114 |
| Arousal Regulatory | -0.19 | [-0.49, 0.11] | 0.208 |

After adjusting for B&B stage and PMI, *PNOC* maintained significant associations with positive valence (β = -0.38, p = 0.035) and arousal regulatory systems (β = -0.43, p = 0.015) (Table 3). *OPRL1* showed no significant associations with RDoC domains in unadjusted models (Table 2), although B&B staging demonstrated significant relationships with cognitive (β = 0.38, p = 0.012) and social domains (β = 0.33, p = 0.026) in the B&B and PMI adjusted models (Table 3).

**Table 3.**
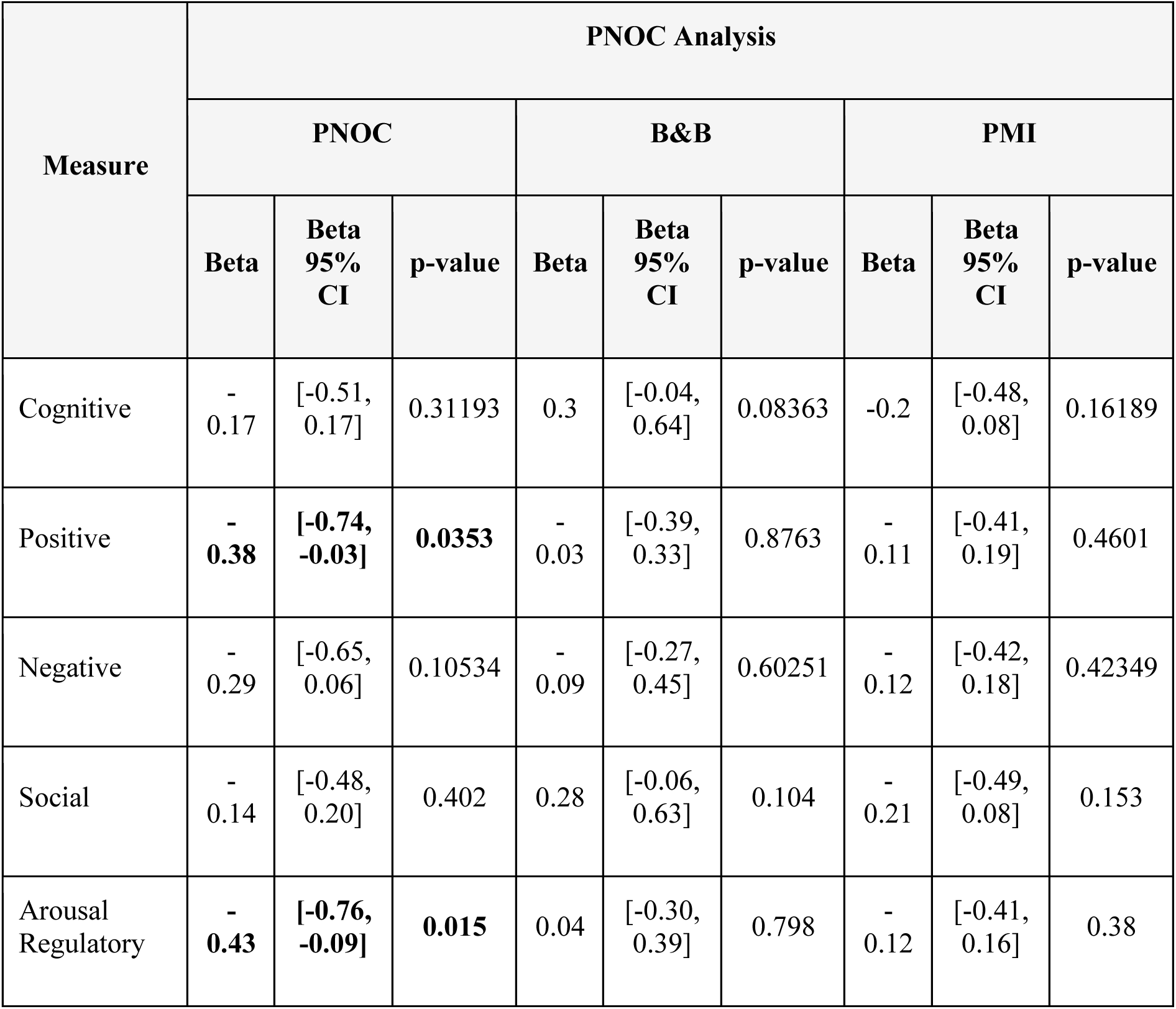
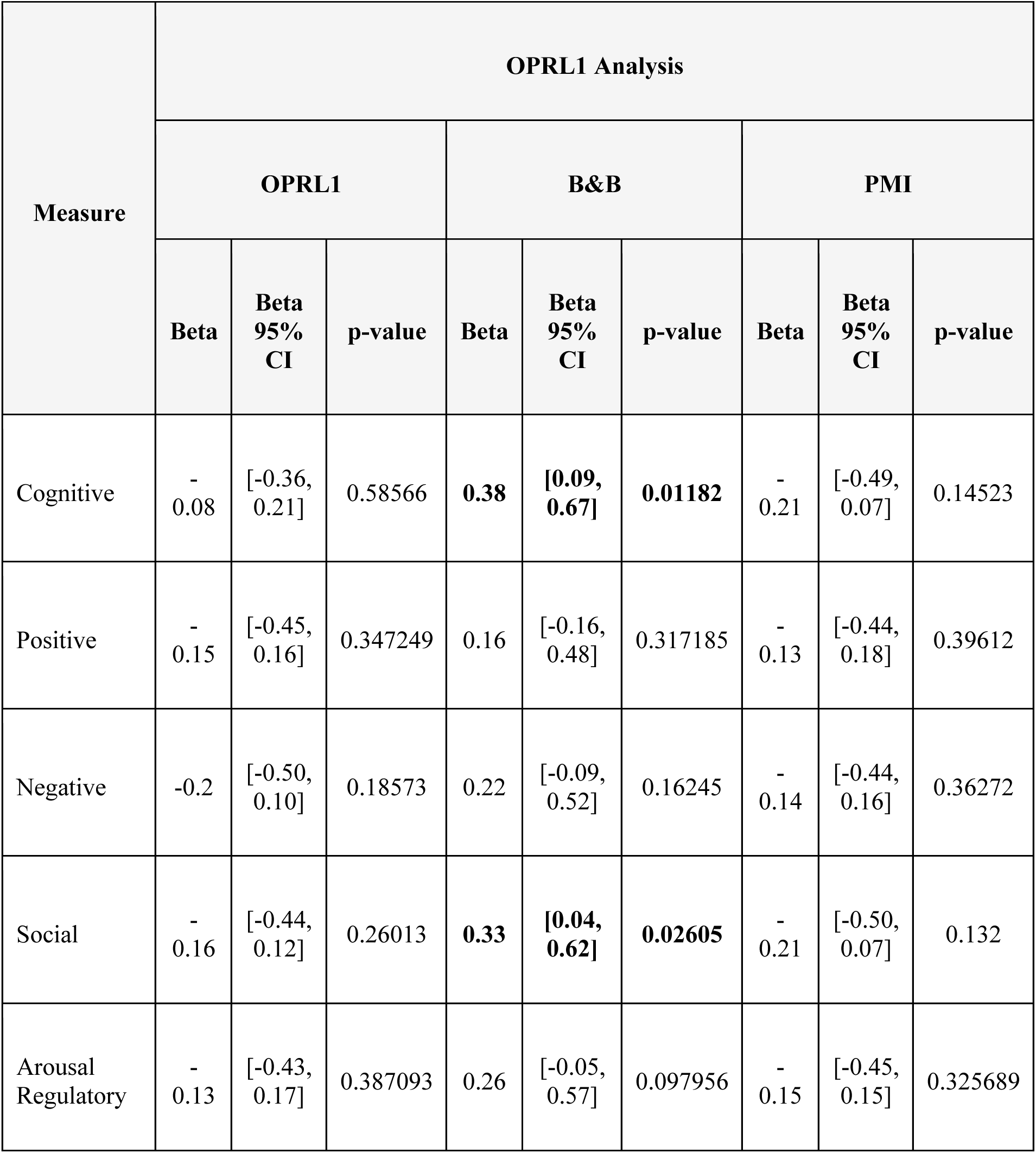
Multi-Covariate Analysis Results.

| Measure | PNOC Analysis |  |  |  |  |  |  |  |  |
| --- | --- | --- | --- | --- | --- | --- | --- | --- | --- |
|  | PNOC |  |  | B&B |  |  | PMI |  |  |
|  | Beta | Beta<br>95%<br>CI | p-value | Beta | Beta<br>95%<br>CI | p-value | Beta | Beta<br>95%<br>CI | p-value |
| Cognitive | -<br>0.17 | [-0.51,<br>0.17] | 0.31193 | 0.3 | [-0.04,<br>0.64] | 0.08363 | -0.2 | [-0.48,<br>0.08] | 0.16189 |
| Positive | -<br>**0.38** | **[-0.74,**<br>**-0.03]** | **0.0353** | -<br>0.03 | [-0.39,<br>0.33] | 0.8763 | -<br>0.11 | [-0.41,<br>0.19] | 0.4601 |
| Negative | -<br>0.29 | [-0.65,<br>0.06] | 0.10534 | -<br>0.09 | [-0.27,<br>0.45] | 0.60251 | -<br>0.12 | [-0.42,<br>0.18] | 0.42349 |
| Social | -<br>0.14 | [-0.48,<br>0.20] | 0.402 | 0.28 | [-0.06,<br>0.63] | 0.104 | -<br>0.21 | [-0.49,<br>0.08] | 0.153 |
| Arousal<br>Regulatory | -<br>**0.43** | **[-0.76,**<br>**-0.09]** | **0.015** | 0.04 | [-0.30,<br>0.39] | 0.798 | -<br>0.12 | [-0.41,<br>0.16] | 0.38 |

| Measure | OPRL1 Analysis |  |  |  |  |  |  |  |  |
| --- | --- | --- | --- | --- | --- | --- | --- | --- | --- |
|  | OPRL1 |  |  | B&B |  |  | PMI |  |  |
|  | Beta | Beta 95% CI | p-value | Beta | Beta 95% CI | p-value | Beta | Beta 95% CI | p-value |
| Cognitive | -0.08 | [-0.36, 0.21] | 0.58566 | **0.38** | **[0.09, 0.67]** | **0.01182** | -0.21 | [-0.49, 0.07] | 0.14523 |
| Positive | -0.15 | [-0.45, 0.16] | 0.347249 | 0.16 | [-0.16, 0.48] | 0.317185 | -0.13 | [-0.44, 0.18] | 0.39612 |
| Negative | -0.2 | [-0.50, 0.10] | 0.18573 | 0.22 | [-0.09, 0.52] | 0.16245 | -0.14 | [-0.44, 0.16] | 0.36272 |
| Social | -0.16 | [-0.44, 0.12] | 0.26013 | **0.33** | **[0.04, 0.62]** | **0.02605** | -0.21 | [-0.50, 0.07] | 0.132 |
| Arousal Regulatory | -0.13 | [-0.43, 0.17] | 0.387093 | 0.26 | [-0.05, 0.57] | 0.097956 | -0.15 | [-0.45, 0.15] | 0.325689 |

### Circadian Rhythmicity Analysis

Investigation of circadian expression patterns revealed distinct temporal regulation of *OPRL1* and *PNOC* (Figure 4, Table 4). In control subjects, *OPRL1* exhibited robust circadian rhythmicity (p = 0.020, R² = 0.37) with an amplitude of 134.33 and phase occurring around Zeitgeber Time 12. This rhythmicity was significantly attenuated in AD cases (p = 0.151, R² = 0.10, amplitude = 71.79), indicating disrupted circadian regulation. The disruption manifested as both reduced amplitude (47% decrease) and loss of consistent 24-hour periodicity in AD cases.

**Figure 4.**
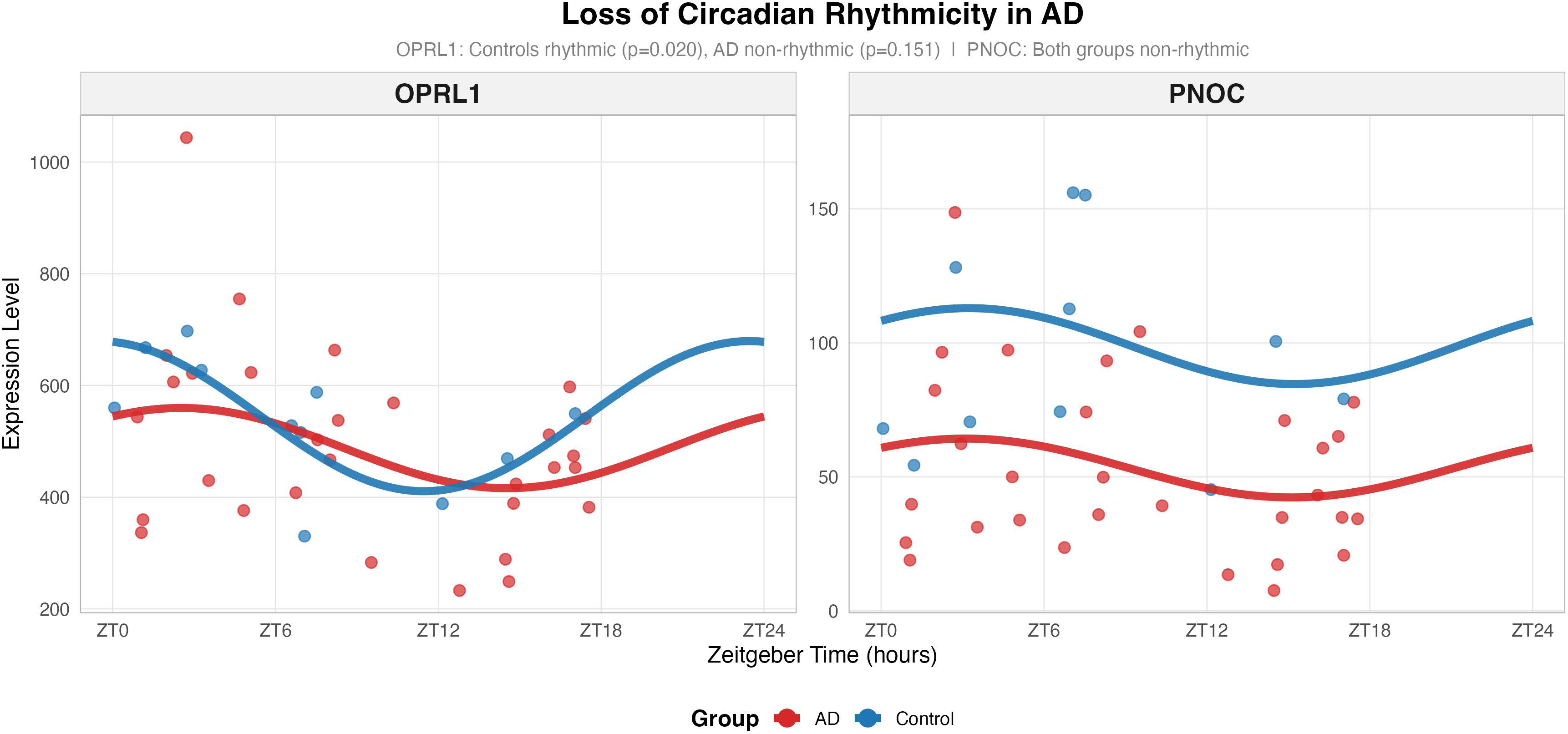
Loss of Circadian Rhythmicity in Alzheimer’s Disease. Circadian expression patterns over 24-hour zeitgeber time (ZT) for *OPRL1* (left panel) and *PNOC* (right panel). Cosinor regression curves show fitted circadian rhythms for controls (blue) and AD cases (red). *OPRL1* exhibits robust circadian rhythmicity in controls (p = 0.020, R² = 0.37) with peak expression around ZT12, which is significantly disrupted in AD (p = 0.151, R² = 0.10). *PNOC* shows minimal circadian variation in both groups. Data points represent individual samples; curves show fitted cosinor models.

**Table 4.**
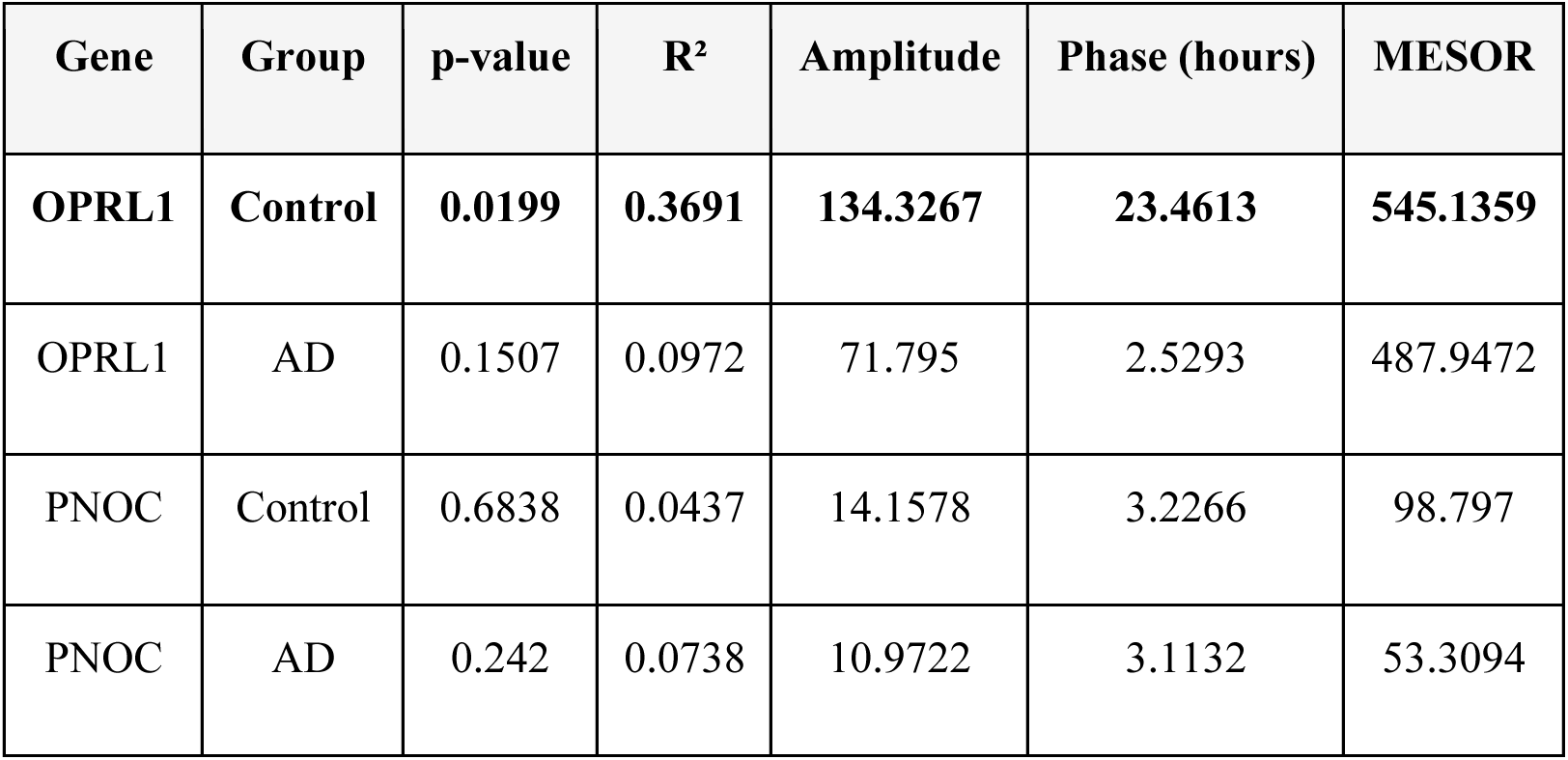
Circadian Rhythm Analysis Results.

In contrast, *PNOC* showed minimal circadian variation in both control (p = 0.684, R² = 0.04) and AD groups (p = 0.242, R² = 0.07), with low amplitude oscillations (A = 14.16 and 10.97, respectively) suggesting that circadian disruption is specific to *OPRL1* regulation in the disease state.

## DISCUSSION

Our findings reveal distinct roles for *PNOC* and *OPRL1* in AD pathology, with implications for both psychiatric symptoms and circadian regulation. The significant associations between *PNOC* expression and specific RDoC domains, particularly positive valence and arousal systems, suggest a mechanistic link between nociceptin signaling and psychiatric manifestations in AD. These relationships persisted after controlling for disease stage and post-mortem interval, indicating that *PNOC* dysfunction may contribute to psychiatric symptoms independently of classical AD pathology.

This study demonstrates a novel approach for in-depth, dimensional analysis of molecular and translational research. While the majority of studies, particularly using post-mortem approaches, focus on classical case control designs, we show that novel NLP-based approaches can be used to bridge clinical gaps. Particularly for chronic and progressive neuropsychiatric disorders and conditions with heterogeneous presentations, symptoms need to be examined in a broader, transdiagnostic and dimensional manner. We studied a neuropathological dimensional cohort across different stages of B&B, from controls (B&B 0-II) to mild and severe AD neuropathology (B&B III-VI).

The strong correlations between *PNOC* expression and RDoC domains provide new insight into the molecular basis of psychiatric symptoms in AD. We observed lower *PNOC* expression in AD patients, consistent with disease-related downregulation. Across subjects, lower *PNOC* expression was associated with higher scores in arousal regulation and positive valence domains. Given that higher arousal regulatory scores reflect greater symptom severity, this finding supports a potential link between reduced *PNOC* expression and dysregulation of arousal systems in AD. The association with positive valence is complex; while moderate PVS scores are typically adaptive, excessively high or dysregulated PVS activation may underlie irritability or affective lability, particularly in neurodegenerative contexts. These findings suggest a role for *PNOC* in modulating both arousal and valence systems, possibly reflecting broader dysfunction in homeostatic regulatory circuits in AD.

The negative association between *PNOC* levels and positive valence scores suggests that reduced nociceptin signaling may contribute to altered reward processing and motivation in AD patients. This finding aligns with previous research demonstrating the role of the nociceptin system in dopaminergic modulation and reward-related behaviors [25,26]. Similarly, the relationship with arousal regulatory systems indicates a potential role in sleep-wake disturbances and energy regulation, which are prominent non-cognitive symptoms in AD that significantly impact patient quality of life and disease progression [27,28].

### Sex-Specific Implications

The observed sex differences in nociceptin system expression align with known sex disparities in AD prevalence and psychiatric symptom profiles. Women comprise approximately two-thirds of AD cases and often exhibit different patterns of neuropsychiatric symptoms compared to men [29]. The higher baseline *PNOC* expression in males (Cohen’s d = -0.482) might contribute to sex-specific vulnerability patterns, particularly given *PNOC*’s association with positive valence and arousal systems in our analyses. Similarly, the sex differences in *OPRL1* expression (Cohen’s d = -0.237) may contribute to differential circadian disruption patterns between sexes in AD.

These findings have important therapeutic implications. The persistence of sex-specific expression patterns even in AD cases suggests that hormonal or genetic factors continue to influence nociceptin system regulation despite disease pathology. Future therapeutic approaches targeting the nociceptin system should consider these sex-specific expression patterns for optimal efficacy, potentially requiring different dosing strategies or treatment timing between men and women.

### Circadian Disruption in AD

The observed disruption of *OPRL1* circadian rhythmicity in AD cases represents a novel finding with potential therapeutic implications. The marked reduction in rhythm amplitude (from 134.33 in controls to 71.79 in AD) and loss of significant rhythmicity (p = 0.151 in AD vs. p = 0.020 in controls) suggests fundamental disruption of circadian regulation. The preservation of *PNOC* rhythmicity indicates that this disruption is specific to the receptor system rather than a global circadian dysfunction.

Several mechanisms might underlie this selective disruption. First, AD pathology may directly affect *OPRL1* transcriptional regulation through disruption of core clock machinery. The suprachiasmatic nucleus, which governs circadian rhythms, shows early pathological changes in AD, potentially disrupting the molecular clock mechanisms that regulate *OPRL1* expression [30]. Second, upstream circadian control elements might be specifically vulnerable to AD-related neurodegeneration.

### Clinical Implications

Our findings present several important implications for AD treatment and research. The early appearance of *PNOC* expression changes suggests potential use as a biomarker for early intervention strategies, particularly given that psychiatric symptoms often precede cognitive decline in AD. Furthermore, the preserved *PNOC* baseline expression suggests that interventions focused on receptor function might be particularly beneficial.

The differential expression patterns between *PNOC* and *OPRL1* suggest distinct therapeutic windows and approaches. The maintenance of higher *OPRL1* expression levels, despite disrupted rhythmicity, suggests that targeting receptor function through chronotherapy or circadian rhythm restoration might be more effective than attempting to modulate ligand levels. Additionally, the distinct roles of *PNOC* and *OPRL1* indicate that combined therapeutic approaches targeting both systems might provide optimal benefits for addressing the multifaceted nature of AD symptomatology.

The sex-specific findings indicate that precision medicine approaches may be necessary when targeting the nociceptin system. Different baseline expression levels between sexes could require sex-stratified clinical trials and potentially different therapeutic doses or administration schedules.

Recent clinical trials with nociceptin receptor antagonists in major depression and substance use disorders provide promising precedents for therapeutic development in AD [31]. The specificity of our findings to particular RDoC domains suggests that such interventions might be particularly beneficial for specific symptom clusters rather than global cognitive decline.

### Future Directions

Further research should focus on several key areas. Longitudinal studies tracking *PNOC*/*OPRL1* changes throughout disease progression will be crucial for establishing causal relationships and determining optimal intervention timing. Additional investigation of the mechanistic interactions between *PNOC*/*OPRL1* and other neurotransmitter systems may reveal new therapeutic targets.

Investigation of the relationship between circadian disruption and cognitive decline progression could inform timing-based therapeutic strategies. Sex-stratified clinical trials will be essential for developing personalized therapeutic approaches. Furthermore, validation of these findings in larger cohorts and different brain regions will be essential for confirming the generalizability of our results.

### Limitations

Several limitations merit consideration in interpreting our findings. The cross-sectional nature of post-mortem studies precludes direct determination of causality between gene expression changes and symptom development. Technical constraints of post-mortem tissue analysis may affect the detection of subtle molecular changes. While our sample size provided adequate statistical power for primary analyses, larger cohorts would be beneficial for investigating sex-stratified subgroup differences and rare variants. Additionally, the NLP algorithms used were developed before the addition of motor systems to the RDoC framework, limiting our analysis to five of the six current domains.

The observed sex differences, while statistically significant and of meaningful effect sizes, should be validated in larger cohorts before being translated into clinical practice. Furthermore, the circadian analysis was based on single time-point measurements, and future studies would benefit from multiple sampling times to better characterize rhythmic patterns.

## CONCLUSIONS

This study provides compelling evidence for distinct roles of *PNOC* and *OPRL1* in AD pathology, with *PNOC* primarily associated with psychiatric symptoms and *OPRL1* showing disrupted circadian regulation. The identification of significant sex differences in both genes highlights the importance of considering biological sex in nociceptin system research and therapeutic development. The findings advance our understanding of the molecular basis of psychiatric symptoms in AD and suggest new therapeutic approaches targeting specific aspects of the nociceptin system through both pharmacological and chronotherapeutic interventions, with consideration for sex-specific treatment strategies.

